# Structural covariance and heritability of the optic tract and primary visual cortex in living human brains

**DOI:** 10.1101/2022.01.27.477973

**Authors:** Toshikazu Miyata, Noah C. Benson, Jonathan Winawer, Hiromasa Takemura

## Abstract

Individual differences among human brains exist at many scales, spanning gene expression, white matter tissue properties, and the size and shape of cortical areas. One notable example is an approximately 3-fold range in the size of human primary visual cortex (V1), a much larger range than is found in overall brain size. A previous study (Andrews et al., 1997) reported a correlation between optic tract cross-section area and V1 size in post-mortem human brains, suggesting that there may be a common developmental mechanism for multiple components of the visual pathways. We evaluated the relationship between properties of the optic tract and V1 in a much larger sample of living human brains by analyzing the Human Connectome Project 7 Tesla Retinotopy Dataset. This dataset includes retinotopic maps measured with functional MRI (fMRI) and fiber tract data measured with diffusion MRI (dMRI). We found a negative correlation between optic tract fractional anisotropy and V1 surface area (*r* = -0.2). This correlation, though small, was consistent across multiple dMRI datasets differing in acquisition parameters. Further, we found that both V1 size and optic tract properties were correlated among twins, with higher correlations for monozygotic than dizygotic twins, indicating a high degree of heritability for both properties. Together, these results demonstrate covariation across individuals in properties of the retina (optic tract) and cortex (V1) and show that each is influenced by genetic factors.

**Significance statement:** The size of human primary visual cortex (V1) has large inter-individual differences. These differences cannot be explained by differences in overall brain size. A previous post-mortem study reported a correlation between the size of the human optic tract and V1. In this study, we evaluated the relationship between the optic tract and V1 in living humans by analyzing a neuroimaging dataset that included functional and diffusion MRI data. We found a small, but robust correlation between optic tract tissue properties and V1 size, supporting the existence of structural covariance between the optic tract and V1 in living humans. The results suggest that characteristics of retinal ganglion cells, reflected in optic tract measurements, are related to individual differences in human V1.

## Introduction

Human brains differ on many levels, including gene expression, morphology of macroscale anatomical landmarks, tissue properties of cortical areas and white matter tracts, size of cortical areas, and perceptual performance. One notable example at the cm scale is primary visual cortex (V1), which varies in size between human brains by up to a factor of 3-4 (Stensaas et al., 1974; Andrews et al., 1997; Amunts et al., 2000; Dougherty et al., 2003; Schwarzkopf et al., 2011; Benson et al., 2021). These individual differences are not explained by variation in whole brain size (Andrews et al., 1997; Benson et al., 2021). Moreover, some studies have found that individual differences in V1 size can be related to properties of visual perception (Duncan and Boynton, 2003; Schwarzkopf et al., 2011; Song et al., 2013, 2015; Genç et al., 2015; Bergmann et al., 2016; Himmelberg et al., 2021). Understanding what factors influence individual differences in V1 size is likely to clarify our understanding of how human visual processing and perception vary across people.

One approach to improve understanding of individual differences is to measure covariance across brain regions (Dougherty et al., 2003; Mechelli et al., 2005). The logic is that if properties of two related structures covary, then these structures may mature by common mechanisms. Andrews and colleagues (1997) used port-mortem human brains to investigate the relationship between V1 and the optic tract (OT), a white matter tract composed of axons from retinal ganglion cells. They found that V1 size was significantly correlated with OT size, suggesting that there are common factors determining individual variability in this fiber tract and V1. This study measured a relatively small number of post-mortem brains. Here we investigated similar questions in a much larger sample of living brains.

Recent progress in non-invasive neuroimaging methods have opened an avenue to quantify the structural properties of cortical areas and white matter tracts in living humans. Specifically, functional MRI (fMRI) enables the identification of visual field maps including V1 (DeYoe et al., 1994; Engel et al., 1994; Sereno et al., 1995; Dumoulin and Wandell, 2008; Wandell and Winawer, 2011), whereas diffusion MRI (dMRI) and tractography enable one to measure properties of white matter tracts including the OT (Conturo et al., 1999; Mori et al., 1999; Sherbondy et al., 2008; Wandell, 2016; Rokem et al., 2017). By taking advantage of these two methods, it is possible to build a large neuroimaging dataset including both fMRI and dMRI data, thus enabling comparisons between them in a large sample (Van Essen et al., 2012; Miller et al., 2016). One such dataset is the Human Connectome Project (HCP) 7 Tesla Retinotopy dataset, which includes fMRI data on retinotopic mapping acquired at 7T and structural and diffusion MRI data acquired at 3T in 178 healthy young adults (Benson et al., 2018). The other unique feature of this dataset is that it includes data acquired from monozygotic (MZ) and dizygotic (DZ) twin pairs, enabling the assessment of heritability.

In this study, we analyzed the HCP 7T Retinotopy dataset to investigate the structural covariance of V1 and OT in living humans. We first identified V1 in all subjects by combining fMRI data with a retinotopic prior based on structural MRI data (Benson and Winawer, 2018) and then quantified its surface area. We then identified the OT in each subject by analyzing dMRI data using tractography (Sherbondy et al., 2008). Finally, we evaluated structural covariance between V1 size and several measures of OT properties. Finally, we examined the heritability of each structure.

## Material and Methods

### Subjects

We analyzed data from 178 subjects (aged 22-35; 107 females, 71 males) whose structural MRI and dMRI data, as well as their retinotopic mapping fMRI data, were collected as part of the Young Adult HCP (Van Essen et al., 2013). Population receptive field models from the retinotopic mapping experiment were solved and published as a separate dataset (Van Essen et al., 2013; Benson et al., 2018). This dataset includes 53 MZ pairs and 31 DZ pairs. All subjects in the HCP dataset provided written informed consent (Van Essen et al., 2013). Further details of the dataset are described in previous publications (Van Essen et al., 2013; Benson et al., 2018).

### MRI data acquisition and preprocessing methods

#### Structural MRI data acquisition and preprocessing

T1-weighted (T1w) structural MR images with an isotropic resolution of 0.7-mm were acquired from a 3T MRI scanner and used for surface-based analysis of fMRI data as well as for identifying V1 in individual subjects. The dataset was preprocessed by the HCP consortium. This preprocessing included automated tissue segmentation implemented in FreeSurfer (Fischl, 2012) and reconstructing the white and pial cortical surfaces (Glasser et al., 2013).

#### Diffusion MRI data acquisition and preprocessing

We analyzed the dMRI dataset acquired by the HCP consortium. In brief, whole-brain dMRI data were corrected at an isotropic resolution of 1.25-mm using 3T MRI. The dMRI data was preprocessed by the HCP consortium to correct eddy-current artifacts and susceptibility-induced image distortions (Andersson et al., 2003; Glasser et al., 2013; Andersson and Sotiropoulos, 2016). The dMRI data consists of three types of diffusion-weighted images (b=1000, 2000, 3000 s/mm^2^) as well as non-diffusion weighted images (b=0 s/mm^2^).

##### Tensor-based analysis

We divided the dMRI data into three datasets acquired with different b-values. We used b=2000 s/mm^2^ as the test dataset since it has a higher signal-to-noise ratio and image contrast, while data acquired with other b-values were used as validation datasets to assess generalizability. We fit a tensor model to each voxel in the dMRI data using a least-squares algorithm implemented in mrDiffusion of vistasoft software (https://github.com/vistalab/vistasoft/). The tensor fits were then used to derive Fractional Anisotropy (FA; Basser and Pierpaoli, 1996).

##### NODDI analysis

We used neurite density and orientation dispersion imaging (NODDI; Zhang et al., 2012) to evaluate tissue properties of the OT. We fit NODDI to the whole dMRI dataset including all b-values using the NODDI MATLAB toolbox (http://mig.cs.ucl.ac.uk/index.php?n=Tutorial.NODDImatlab). From these fits, we obtained intra-cellular volume fractions (ICVF) and the orientation dispersion index (ODI).

#### fMRI Image acquisition and preprocessing

Full details of the fMRI data acquisition and preprocessing steps are provided in previous publications (Glasser et al., 2013; Vu et al., 2017; Benson et al., 2018). In brief, wholebrain fMRI data were collected at an isotropic resolution of 1.6-mm using 7T MRI. During the fMRI data acquisition, subjects were instructed to perform a fixation task that required them to maintain gaze at the center of the screen; simultaneously, they were presented with retinotopic mapping stimuli. These stimuli were constructed from slowly moving apertures that contained dynamic colorful textures (Dumoulin and Wandell, 2008; Benson et al., 2018). FMRI data were preprocessed with the HCP pipelines (Glasser et al., 2013).

### MRI data analysis methods

#### Diffusion MRI data analysis

##### Defining ROIs for tractography

We identified regions of interest (ROIs) for tractography based on T1w images in each individual subject, using the same method employed in previous work (Ogawa et al., 2014; Takemura et al., 2019). Briefly, the optic chiasm was defined based on Freesurfer’s automated segmentation (Fischl, 2012). We then defined the lateral geniculate nucleus (LGN) ROIs manually using a previously-validated method (Takemura et al., 2019). This method identifies the approximate location of the LGN by following streamlines found using deterministic tractography from a seed-region in the optic chiasm to their termination points. The LGN ROI is defined as a 4-mm radius sphere covering these endpoints (Takemura et al., 2019). We also identified the V1 ROI for tractography using the Brodmann Area atlas implemented in Freesurfer. We used this atlas, rather than V1 boundaries identified using fMRI data because, for the purpose of tractography, the larger Brodmann V1 ROI may improve the sensitivity of tractography for the purposes of identifying the optic radiation. The V1 ROIs were redefined using functional data for the purpose of quantifying V1 size (see below).

##### Tractography

We performed tractography on the dMRI data (with b=2000 s/mm^2^) to identify the OT using ConTrack (Sherbondy et al., 2008). ConTrack is a probabilistic tractography method for identifying the most probable path of the white matter tract connecting two ROIs. Specifically, we sampled 5,000 candidate streamlines connecting the optic chiasm and the LGN ROIs in both hemispheres (angle threshold, 90°; step size, 1 mm; maximum streamline length, 80 mm). We then refined OT streamlines using the following criteria. First, we selected 100 streamlines with the highest scores in the ConTrack scoring process (Sherbondy et al., 2008). Second, we removed streamlines with (1) length > 3 S.D. longer than the median streamline length in the tract, or (2) position > 3 S.D. away from the median position of the tract using AFQ MATLAB toolbox (Yeatman et al., 2012).

In addition to the OT, we also identified the optic radiation. For the optic radiation, we used ConTrack to generate streamlines connecting the LGN and V1 ROIs and to reject outlier streamlines. We used the identical procedure for identifying the optic radiation as used in a previous study (Takemura et al., 2019).

##### Estimating the cross-section area of the OT

We quantified the cross-section area of the OT to match the dependent measures from previous anatomical work (Andrews et al., 1997). To do this, we first identified voxels that intersected the OT streamlines in each coronal section of the dMRI data. We then multiplied the voxel size in the section (1.25 x 1.25 mm^2^) by the voxel count to calculate the cross-section area in each coronal section. Finally, we averaged the cross-section area across coronal sections to obtain the mean cross-section area of the OT in each individual subject. When averaging, we excluded the 10% of sections nearest the optic chiasm ROI and the 10% of sections nearest the LGN ROI. This reduces the possibility of including the optic chiasm or the LGN in our estimate of OT cross-section area. For comparisons with V1 data, we averaged the data from the left and right hemisphere of each individual subject.

##### Evaluating tissue properties of the OT

We evaluated tissue properties of the OT based on methods from previous work using the AFQ MATLAB toolbox (Yeatman et al., 2012; Duan et al., 2015; Takemura et al., 2019). Briefly, we resampled each streamline to 100 equidistant nodes. Tissue properties (FA, ICVF, and ODI) were calculated at each node of each streamline. The properties at each node were summarized by taking a weighted average of the tissue measurements of each streamline within that node. The weight of each streamline was based on the Mahalanobis distance from the tract core. We excluded the first and last 10 nodes from the calculation of the tissue property of the tract core to minimize any partial voluming effects with neighboring structures, which often occur near the endpoints of the tract. We averaged data from the remaining 80 nodes to obtain a single-number summary of each tissue property (FA, ICVF, and ODI) for each subject. The data from the left and right hemisphere were averaged for comparisons with V1. We performed the same analysis for the optic radiation.

#### Functional MRI data analysis

##### Bayesian retinotopy analysis for identifying the V1 surface area

We measured the surface area of V1 in each subject by analyzing fMRI and structural MRI data. To do so, we first performed Bayesian analysis of retinotopic maps as described in previous work (Benson and Winawer, 2018). This method combines traditional fMRI-based retinotopic mapping (Dumoulin and Wandell, 2008) with a retinotopic template (Benson et al., 2012, 2014). Specifically, this method identifies the retinotopic parameters of each surface vertex in visual cortex, as well as the boundaries between maps, using Bayesian inference in which structural MRI acts as a prior constraint and fMRI data as an observation. This analysis procedure has been demonstrated to reliably identify properties of the early retinotopic areas (V1/V2/V3) and is implemented in the publicly available neuropythy library (https://github.com/noahbenson/neuropythy).

The V1 ROIs were projected onto the mid-gray cortical surface mesh for measuring surface area. Surface area for each subject’s V1 was calculated by summing the area of the mesh triangles and partial sub-triangles they contained. We averaged the surface areas of V1 across hemispheres for a comparison with the OT data.

### Experimental design and statistical analyses

#### Statistical evaluation

##### Correlation between the OT and V1

We quantified the correlation between the OT measurements (cross-section area, FA, ICVF and ODI) and V1 surface area by calculating the Pearson correlation coefficient between them. We report the P-value of the Pearson correlation and define the statistical significance (α) as *P* = 0.0125, which is equivalent to *P* = 0.05 after Bonferroni correction for four comparisons. We note that this threshold might be too stringent for considering a correlation between FA and ODI (Zhang et al., 2012).

##### Evaluating inter-hemispheric correlation

We also tested the inter-hemispheric correlation of the OT measurements for evaluating the reliability of the measurement. To this end, we calculated the Pearson coefficient across hemispheres. We test interhemispheric correlations as a proxy for test-retest reliability on the assumption that the true values (without measurement noise) are highly correlated between hemispheres.

##### Evaluating the influence of the inclusion of twin pairs on the correlation

We evaluated how much the inclusion of twin pairs, which may not be fully independent samples, affects the correlation between OT and V1. To perform this analysis, we generated a subsample of subjects in which we removed one member of each twin pair from the whole dataset and reassessed the OT-V1 correlation, leaving 94 subjects. Because either of the members of each twin pair could be removed, there are many to select the subset of subjects. Hence we generated subsamples of 94 subjects 10,000 times, and each time calculated the OT-V1 correlation to obtain a distribution of the correlation coefficient after the removal of twin pairs. This distribution was compared with the null distribution of the correlation coefficient, which was obtained by randomly choosing 94 subjects 10,000 times (without regard to twin status) and shuffling the association between OT and V1 data (see Supplementary Figure 3).

#### Evaluating heritability

We evaluated the heritability—i.e., the fraction of the variance for a given trait that is attributable to genetics—of the OT FA and of the surface area of V1 by comparing the correlations of these measurements between MZ twin-pairs to the correlations between DZ twin-pairs. We used the intraclass correlation coefficient (ICC; Shrout and Fleiss, 1979) when examining twin-pairs. We then employed Falconer’s formula (Falconer and Mackay, 1996) to estimate OT FA and V1 surface area heritabilities.

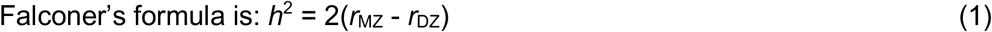

where *h*^2^ is the heritability index and *r*_MZ_ and *r*_DZ_ are the ICCs between MZ pairs and DZ pairs, respectively.

### Code Accessibility

Code for reproducing figures and statistical analyses of this work will be publicly available at a public repository ([URL will be inserted upon an acceptance of this manuscript]). Codes were written in MATLAB 2015a and tested on Ubuntu 14.04 LTS. However, we cannot release data for reproducing the twin analyses because the family structure data belongs to the HCP’s restricted dataset and thus cannot be shared without consent of the HCP Consortium. To reproduce analysis requiring family structure, researchers must apply for access to “Restricted Data” on ConnectomeDB (see https://www.humanconnectome.org/study/hcp-young-adult/document/wu-minn-hcp-consortium-restricted-data-use-terms).

## Results

We identified V1 and the OT by analyzing functional, structural, and diffusion MRI datasets in the HCP Young Adult and the HCP 7T Retinotopy datasets (Figure 1A and 1B), in 178 subjects. Similar to previous work (Stensaas et al., 1974; Andrews et al., 1997; Amunts et al., 2000; Dougherty et al., 2003; Schwarzkopf et al., 2011), and as reported recently for this dataset (Benson et al., 2020), there is a considerable degree of inter-individual difference in V1 surface area, spanning a two-fold range (Figure 1C). Cross-section area and FA of the OT also exhibited considerable inter-individual differences (Figure 1D and 1E). In subsequent analyses, we focused on the relationship between individual differences of V1 and the OT in the HCP 7T Retinotopy dataset.

**Figure 1.**
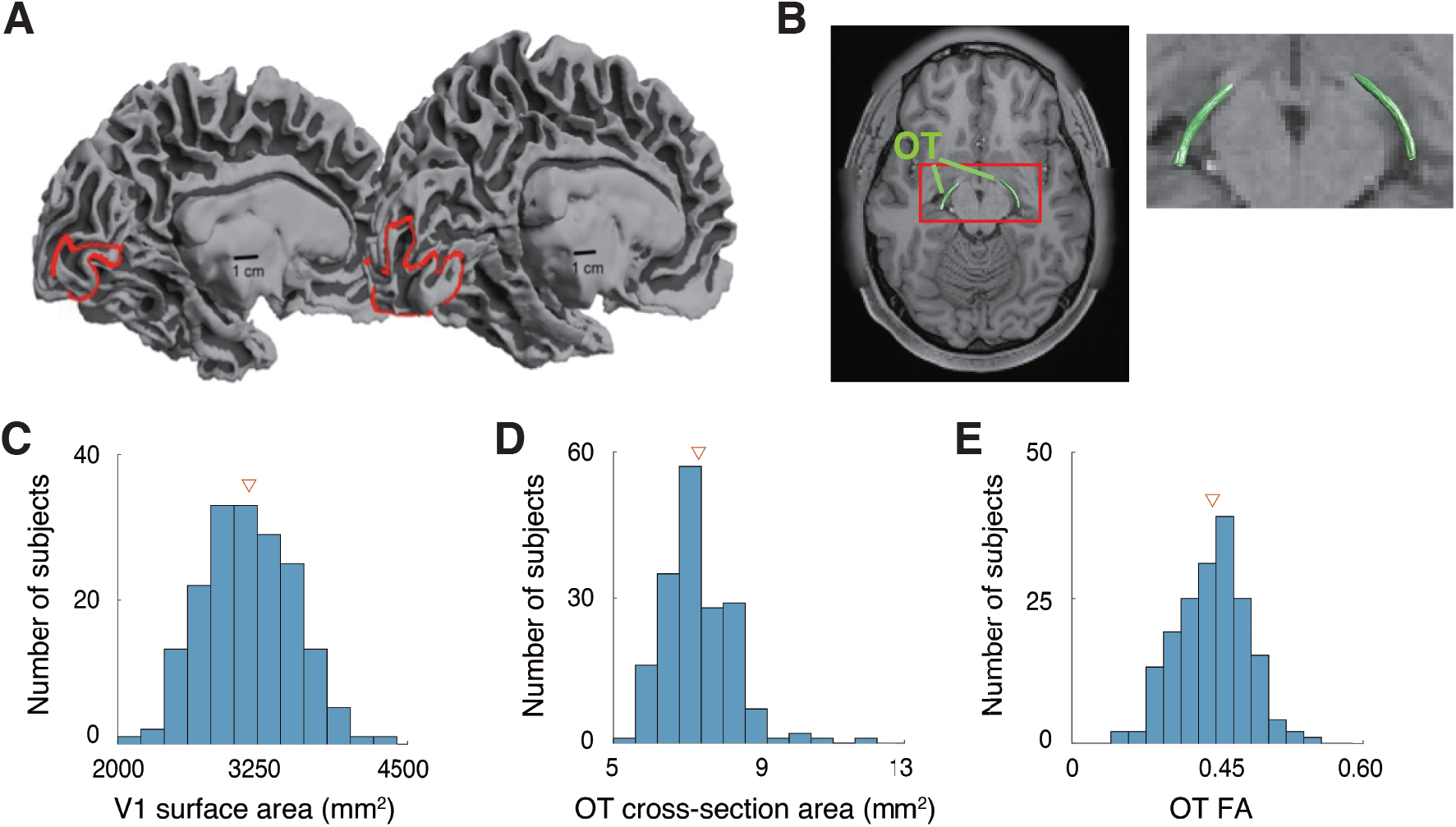
Individual variability of structural properties of the early visual system in human brains. **A.** Human V1 identified by fMRI-based retinotopy measurements. The locations of V1 in the left hemisphere of two subjects are shown as red lines on the mid-gray surface. The V1 surface area of these two hemispheres differs by a factor of 3.5 (left, subject 958976, 620 mm^2^; right, subject 100610, 2180 mm^2^). This image is adapted from Benson et al. (2020). **B.** Human optic tract (OT) of a representative subject (subject 100610), identified by dMRI-based tractography, overlaid on an axial slice of the subject’s T1-weighted image. The red rectangle on the left panel indicates the region that is magnified on the right. **C-E.** Histograms of the V1 surface area (**C**), OT cross-section area (**D**), and OT FA (**E**) of individual subjects (N = 178) averaged across the left and right hemispheres. Triangles in each plot depict mean across subjects.

### Correlation between V1 surface area and OT cross-section area

We first examined the correlation between V1 surface area and OT cross-section area, since such a correlation was reported in post-mortem anatomical work (Andrews et al., 1997). Figure 2 depicts a scatter plot comparing the V1 surface area and the OT cross-section area. We did not find a statistically significant correlation between these measurements (*r* = 0.05; *P* = 0.5). Therefore, we failed to replicate the previous anatomical finding in an *in vivo* neuroimaging dataset.

**Figure 2.**
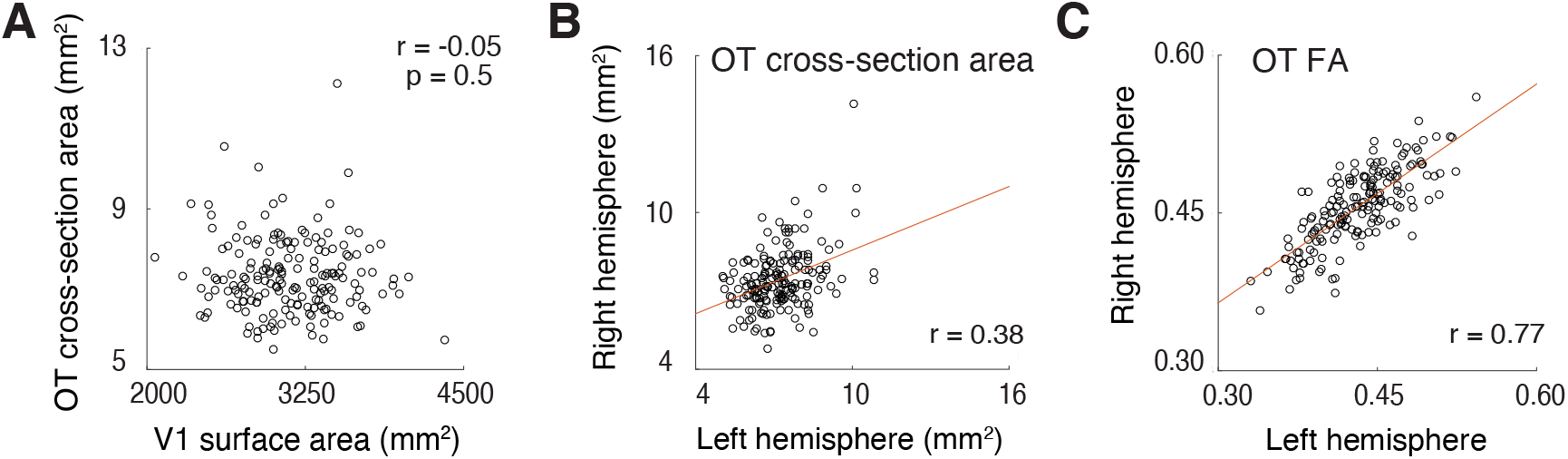
**A.** No significant correlation between V1 surface area (horizontal axis) and OT cross-section area (vertical axis). **B-C.** Inter-hemispheric correlation of OT cross-section area (**B**) and OT FA (**C**). The OT crosssection area showed smaller inter-hemispheric correlation (*r* = 0.38) than OT FA (*r* = 0.77), presumably because of instability of dMRI-based measurements on the OT size.

One possible interpretation of this negative result is that there is a high degree of measurement noise in our OT cross-section area estimates due to low spatial resolution. The voxel size of the dMRI dataset in the 3T HCP Young Adult dataset is 1.25 mm, isotropic. The cross-section area of the OT ranges from 5.1 mm^2^ to 12.1 mm^2^, according to Andrews et al. (1997). If we aim to estimate the volume of such a small tract with 1.25 mm voxel size, a large proportion of the OT voxels will be located on the border between the OT and neighboring tissue, and thus the estimates of OT size will depend on the placement of voxels (Supplementary Figure 1A). This will cause additional noise in the measurements and will mask true inter-individual differences in OT size. In contrast, measurements of diffusivity properties, such as FA, are likely to be less prone to this issue since one can calculate a weighted summary of tract FA by minimizing the weight of voxels located far from the tract core (Yeatman et al., 2012).

One metric of reliability is inter-hemisphere correlations. We computed this for the OT cross-section area and FA. A key assumption in this analysis is that measurements of the left and right OT in the same subject should be similar. While this assumption is only likely to be approximately correct, it provides a reasonable estimate of reliability since inter-hemisphere correlation was very high in a previous post-mortem study (Supplementary Figure 2, *r* = 0.84, in data presented in Andrews et al., 1997). We found that the correlation between left and right OT cross-section areas was much smaller (*r* = 0.38) than that between left and right OT FA (*r* = 0.77). This difference in correlation coefficient was statistically significant (*P* < .001). This result suggests that the OT crosssection area measurements based on dMRI are noisier than the FA measurements. This result motivated us to investigate the correlation between V1 surface area and OT FA, since FA is a more robust measurement for characterizing individual differences in OT structural properties.

### Correlation between V1 surface area and OT tissue properties

We investigated the correlation between V1 surface area and OT FA. Figure 3A depicts a scatter plot between these two quantities. We found a small, but statistically significant negative correlation between them (Figure 3A; *r* = −0.19, *P* = 0.01), suggesting the existence of structural covariance between the surface area of V1 and tissue properties of the OT.

**Figure 3.**
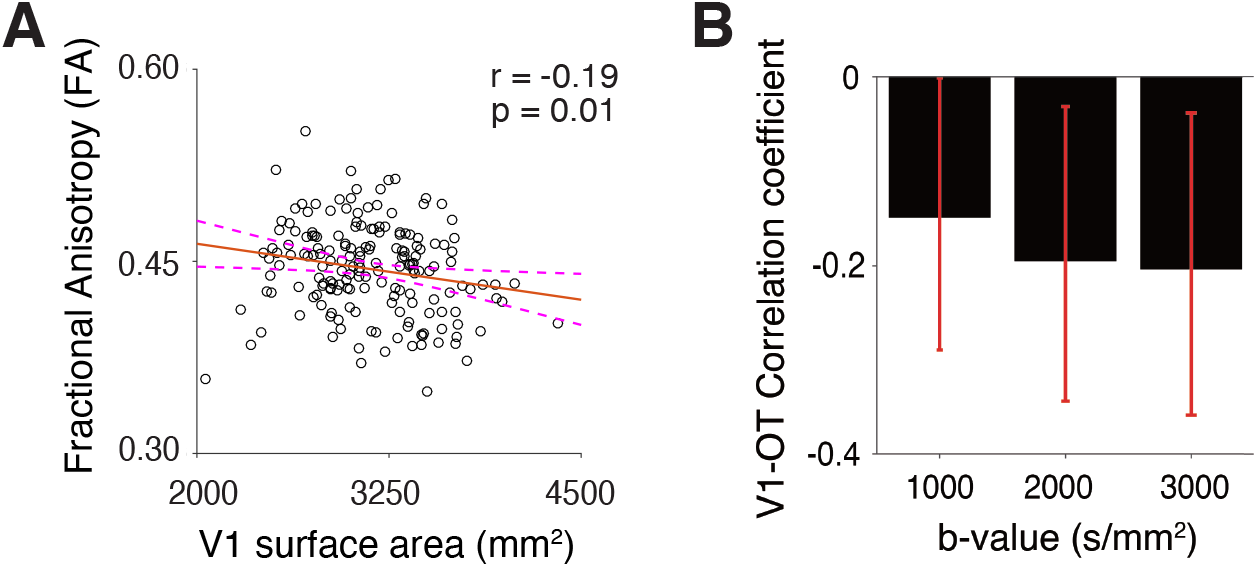
The correlation between V1 surface area and OT tissue property. **A.** A scatter plot of the V1 surface area (horizontal axis) and OT FA (vertical axis). The correlation between V1 surface area and FA along the OT was small, but statistically significant (*r* = −0.19, *P* = 0.009, N = 178). **B.** The correlation between V1 surface area and FA along the OT was replicated in dMRI data acquired with different b-values (b=1000 s/mm^2^, weaker diffusion weighting; b=3000 s/mm^2^, stronger diffusion weighting). The vertical axis depicts the correlation coefficient between V1 surface area and OT FA. We performed the bootstrap 10,000 times for each b-value to compute the 95% confidence interval of this probability. Error bars indicate this interval, estimated by bootstrapping.

We then asked whether this observed correlation in the main analysis, which used dMRI data acquired with b = 2,000 s/mm^2^, generalizes. To this end, we performed the same analyses across different b-values (Lerma-Usabiaga et al., 2019). We found that the negative correlation between V1 surface area and OT FA is similar in other dMRI datasets with different acquisition parameters (Figure 3B; b = 1,000 s/mm^2^, *r* = −0.15; b = 3,000 s/mm^2^, *r* = −0.20). This result suggests that the observed structural covariance between V1 and OT FA is unlikely to be due to measurement noise.

The HCP 7T Retinotopy dataset includes many twin subjects (see Materials and Methods) and twin pair subjects may show similar properties in OT and V1. We evaluated how much the inclusion of twin pairs in the dataset impacted the correlation between OT FA and V1 surface area. First, we randomly sampled 94 subjects from the whole dataset while avoiding the inclusion of twin pairs (for both MZ and DZ twins). We repeated this sampling procedure to generate many subsamples (see Materials and Methods). Then for each subsample we calculated the correlation coefficient between OT FA and V1 surface area (blue in Supplementary Figure 3). The median correlation coefficient among subsamples was −0.21, which is comparable to that obtained in the main analysis of data from all subjects (Figure 3A). The distribution of correlation coefficients among subsamples avoiding twin pairs (blue) significantly differs from the distribution of correlation coefficients when randomly shuffling the association between OT FA and V1 surface area (orange in Supplementary Figure 3). This result suggests that a negative correlation in OT FA and V1 surface area (Figure 3A) is preserved after excluding the impact of twin pairs.

We also investigated whether V1’s observed structural covariance with OT generalized to the other visual white matter tract. To this end, we identified the optic radiation (OR), which is a geniculo-cortical pathway connecting the LGN and V1, from the dMRI dataset (Supplementary Figure 4A). We did not find a significant correlation between OR FA and V1 surface area, unlike with the OT (*r* = 0.04; *P* = 0.61).

### Possible microstructural basis of OT-V1 correlation evaluated by NODDI

While FA is a widely used metric with high reproducibility, it is based on a simplistic diffusion tensor model, and the interpretation of its microstructural origin is inherently ambiguous. To better understand the microstructural basis of the OT-V1 correlation, we used NODDI (Zhang et al., 2012), which is a multi-compartment model of dMRI data. NODDI enables one to estimate ICVF and ODI, which are thought to be correlated with axonal volume fraction and the orientation dispersion of fibers, respectively. Similar to FA, these metrics are strongly correlated between hemispheres (Supplementary Figure 5; ICVF, *r* = 0.80; ODI, *r* = 0.85), suggesting high measurement reliability.

Figure 4 depicts scatter plots comparing V1 surface area and OT ICVF and ODI. We found a small, but statistically significant positive correlation with V1 surface area in both ICVF (Figure 4A; *r* = 0.19, *P* = 0.01) and ODI (Figure 4B; *r* = 0.22, *P* = 0.003). This result suggests that a negative correlation between FA and V1 surface area can be explained by greater axonal density and orientation dispersion in subjects with a larger V1 (see “Microstructural origin of OT-V1 correlation” in the Discussion).

**Figure 4.**
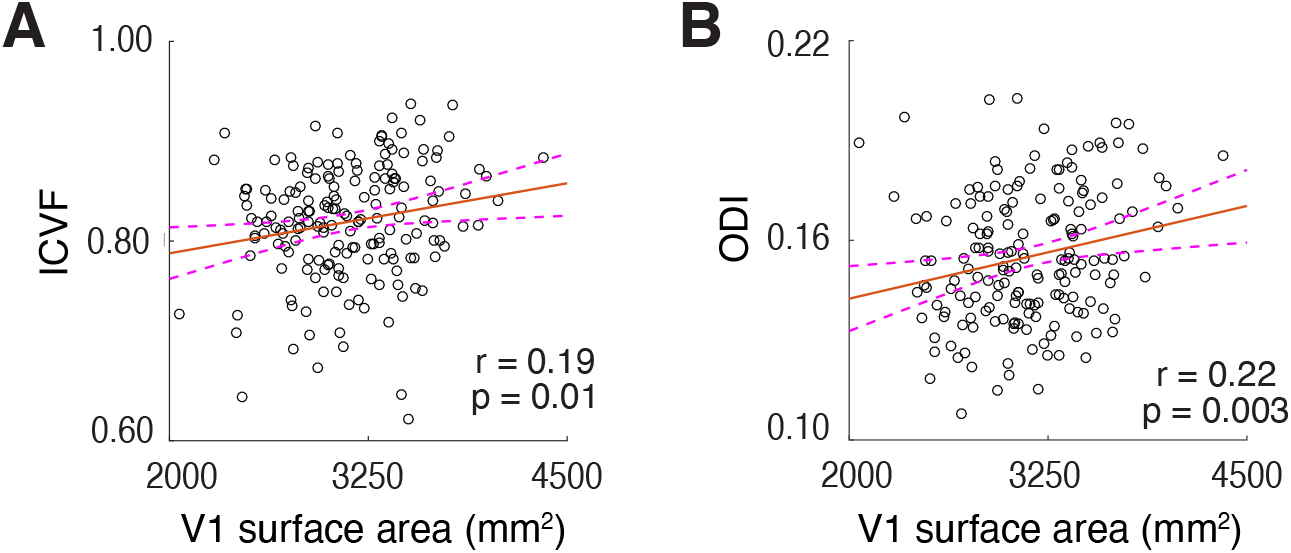
The correlation between V1 surface area and microstructural properties of the OT as quantified by NODDI (**A.** ICVF; **B.** ODI). Other conventions are identical to those used in Figure 3A.

### Heritability of human OT and V1 structural properties

Finally, we evaluated the heritability of OT FA and V1 surface area to understand whether individual variability of these measurements derives from genetic factors. To this end, we evaluated the correlation of OT FA and V1 surface areas between MZ and DZ twin pairs and calculated the heritability index (Falconer and Mackay, 1996). For both V1 surface area and OT FA, correlations between MZ twins were higher than correlations between DZ twins, suggesting a considerable degree of heritability (Figure 5A, Falconer’s h^2^ = 0.58 for OT FA; Figure 5B, Falconer’s h^2^ = 0.28 for V1 surface area; see Supplementary Figure 6 for confidence intervals).

**Figure 5.**
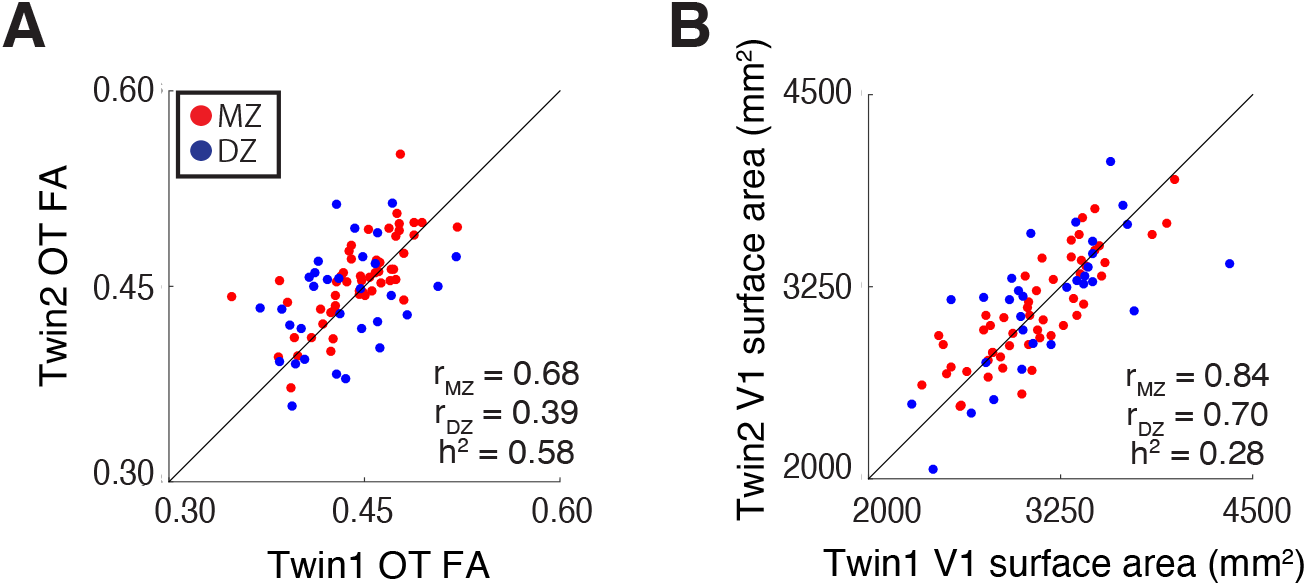
Heritability of structural properties of OT and V1 evaluated by analysis of twin data. The scatter plot depicts a comparison of twin pair data for OT FA (**A**) and V1 surface area (**B**). Red dots depict data from MZ twins whereas blue dots depict data from DZ twins. The black lines depict the line of equality (x = y). The heritability index (h^2^) was calculated by Falconer’s formula of heritability based on ICC in MZ and DZ data (r_MZ_ and r_DZ_).

## Discussion

We evaluated the structural covariance between V1 and OT in living humans, which was hypothesized based on previous anatomical work (Andrews et al., 1997), by analyzing fMRI and dMRI data in the HCP 7T Retinotopy dataset. We did not find a significant correlation between OT size and V1 surface area, presumably due to the instability of the OT size measurements from the dMRI data (Figure 2). In contrast, we found a significant negative correlation between V1 surface area and OT FA, and this correlation generalized across acquisition parameters (Figure 3). Moreover, we found that a negative correlation between OT FA and V1 could be explained by individual differences in intracellular diffusivity and orientation dispersion along the OT (Figure 4). Finally, we found that there is a considerable degree of heritability in both OT FA and V1 surface area (Figure 5), suggesting that genetic factors influence both measurements.

### Structural covariance between OT and V1 in living humans

In contrast to previous work (Andrews et al., 1997), we did not find a statistically significant correlation between OT cross-section area and V1 surface area (Figure 2A). This is likely because the spatial resolution of dMRI data (1.25 mm^3^) is not sufficient to robustly identify the cross-section area of the OT, which ranges from 5.1 to 11.3 mm^2^ in post-mortem living brains (Andrews et al., 1997). If we assume that the OT cross-section area is comparable in the living human brain, the number of voxels covering OT in each coronal section should range from 3 to 7. Because the estimation of OT cross-section area significantly depends on the spatial arrangement of voxels and partial voluming with neighboring tissues (Supplementary Figure 1A), it is likely that measurements of the OT cross-section area are unstable at this spatial resolution. In fact, we also found the interhemispheric correlation of the OT cross-section area measurements from dMRI data to be low (*r* = 0.37; Figure 2B), confirming that this measurement is unstable.

In contrast, FA along the OT can be reliably measured from this dataset because in the AFQ analysis pipelines, as used by Yeatman et al. (2012), the tract profile of FA was calculated by taking a weighted average where the weights were defined based on distance from the tract core (see Materials and Methods). This procedure is known to be highly reliable in terms of test-retest reliability (Kruper et al., 2021) and has successfully identified OT tissue abnormalities in retinal disease patients (Ogawa et al., 2014; Takemura et al., 2019). In fact, our results show that the inter-hemispheric correlation of OT FA is higher than that of OT cross-section area, suggesting that this is a more reliable dMRI-based metric for OT structural properties (Figure 2C). Because OT FA is a reliable measurement and because the significant negative correlation between OT FA and V1 surface area (Figure 3A) has been generalized across acquisition parameters (Figure 3B), our results support the existence of structural covariance between the OT and V1 in living human brains.

### Microstructural origin of OT-V1 correlation

FA is a fairly reproducible dMRI-based metric of white matter microstructure (Kruper et al., 2021), but it does not have a direct correlation with specific microstructural properties, such as the properties of axons and myelin (Wandell and Le, 2017; Assaf et al., 2019). Therefore, it is challenging to interpret the microstructural origin of the OT-V1 correlation solely from FA results (Figure 3). One plausible hypothesis is that individuals with larger V1s have more retinal ganglion cells (RGCs), and therefore those individuals will have more neurons in V1, which will result in greater V1 surface area. In fact, this hypothesis is supported by a correspondence of radial field asymmetries between retina and cortex at the population level (Kupers et al., 2021). If this is the case, the OT-V1 correlation may suggest that OT microstructural properties measured by dMRI are related to the number of RGCs. This interpretation remains speculative at this point, since the HCP 7T Retinotopy dataset does not have a direct measurement of retinal ganglion cells.

One might wonder why the correlation between OT FA and V1 is negative, rather than positive, if the basis of individual differences are derived from differences in numbers of RGCs. To clarify this point, we used NODDI (Zhang et al., 2012), which is a multi-compartment model providing ICVF and ODI, two properties that are hypothesized to be correlated with axonal volume and orientation dispersion, respectively (Mollink et al., 2017; Schilling et al., 2017). We found that both ICVF and ODI along the OT showed a positive correlation with V1 surface area (Figure 4). This result is consistent with the interpretation that individuals with larger V1 have larger axonal volume with more dispersed fiber orientations along the OT. This suggests that individuals with more RGCs must have more axons (resulting in a larger ICVF), but the configuration of axonal orientation in such individuals may also be more dispersed than that of individuals with fewer RGCs (Supplementary Figure 1B). Regardless, this hypothesis requires further evaluation by anatomical studies, since the microstructural interpretation of NODDI has some degree of uncertainty (Jelescu et al., 2016). It is also known that there is substantial individual variation in acuity at the fovea and in cone density at the fovea (Curcio et al., 1987). An important open question is whether differences in OT properties and V1 surface area can be traced all the way back to the photoreceptor mosaic.

### No evidence of correlation between OR and V1

We did not find a significant correlation between OR FA and V1 surface area (Supplementary Figure 4B). This is counter-intuitive, since the OR includes axons that directly project to V1. We speculate that this lack of significance can be explained by the fact that the OR comprises heterogenous fiber populations. For example, it is known that the OR includes feedback axons from V1 to LGN (Ichida and Casagrande, 2002; Angelucci and Sainsbury, 2006). In addition, a recent anatomical study reported that axons from the pulvinar merge into the OR and follow a path similar to that of axons from the LGN (Takemura et al., 2020). Therefore, unlike the OT, in which all axons are feedforward axons from RGCs, feedforward axons can only explain a part of the variance in dMRI measurements along the OR. Resolving this uncertainty requires further anatomical investigations into how different axonal populations are spatially organized within the OR.

### Possible underlying mechanism of structural covariance between OT and V1

While it is difficult to identify the underlying mechanisms of the structural covariance between OT and V1, there are, at minimum, several possible interpretations. First, since the OT does not contain feedback axons from V1, it is natural to infer that the OT influences V1, but that V1 would not conversely influence the OT. Accordingly, one hypothesis is that the development of the OT affects the maturation of V1, resulting in a correlation between them. The second possibility is that a common genetic factor affects the development of both OT and V1, resulting in structural covariance, without a direct relationship between the OT and V1.

Although these two possibilities are not mutually exclusive, we investigated the twin data included in the HCP 7T Retinotopy Dataset to better interpret the structural covariance of the OT and V1. We found that MZ twin pairs had higher correlations than DZ twins (Figure 5) for both the OT FA and V1 surface area, suggesting a considerable degree of heritability in these measurements. While the specific genetic factors contributing to the structural covariance between the OT and V1 remain uncertain, these results suggest that one plausible source of that covariance is common genetic factors. An extension of this study, by combining developmental neuroimaging and transcriptomics (Natu et al., 2021), may provide a more precise understanding of the developmental and genetic mechanisms of the structural covariance between the OT and V1.

In conclusion, we found a small, but statistically significant correlation between OT microstructural properties and V1 surface area. This correlation was generalized across datasets acquired with different parameters. These results support the existence of structural covariance between the OT and V1, as hypothesized from previous anatomical work (Andrews et al., 1997). Since both the OT and V1 have a considerable degree of heritability, one plausible source of the structural covariance might be common genetic factors.

## Acknowledgments

We thank Yusuke Sakai in support of data curation and analysis. This study was supported by the Japan Society for the Promotion of Science (JSPS) KAKENHI (JP17H04684 and JP21H03789 to H.T.), National institute of Information and Communications Technology project for Collaborative Research in Computational Neuroscience (CRCNS): Innovative Approaches to Science and Engineering Research on Brain Function (to H.T.), and National Eye Institute CRCNS: Innovative Approaches to Science and Engineering Research (to N.C.B. and J.W.).

## Extended data of

**Supplementary Figure 1.**
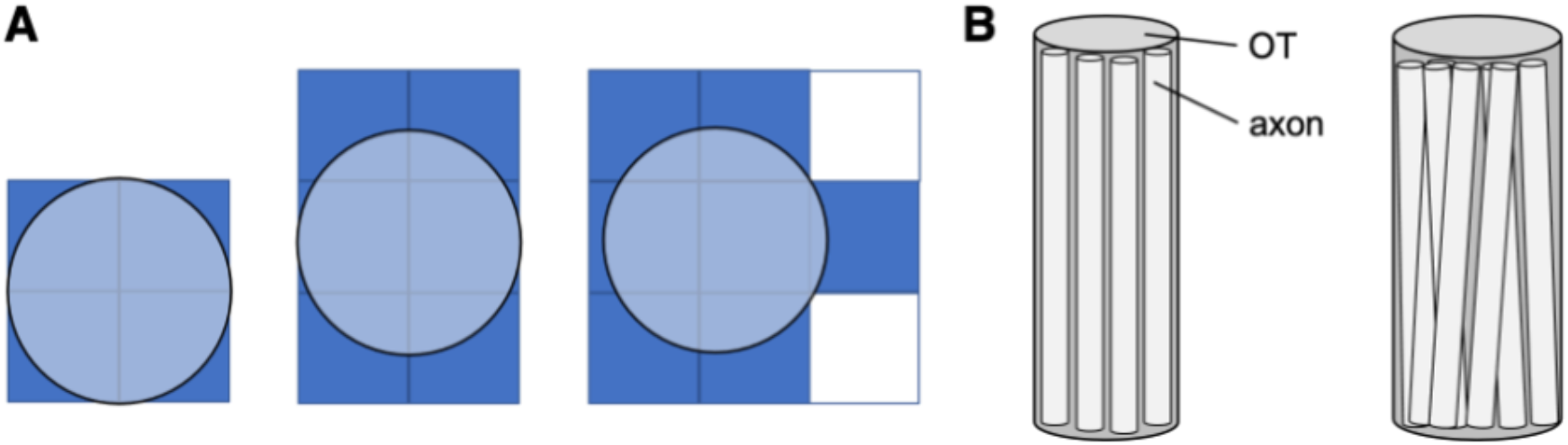
**A.** Schematic figure explaining how the spatial arrangement of voxels impacts the estimate of optic tract (OT) cross-section area when the spatial resolution of the measurements is limited. In all panels, the true cross-section area of the OT (light blue circle) is identical. However, the voxel count (dark blue) significantly differs across these three cases due to differences in the placement of the voxels (squares). **B.** Schematic figure explaining how the number and the spatial configuration of axons correlates with primary visual cortex (V1) surface area. The pattern of OT-V1 correlation (Figure 4) raises the hypothesis that individuals with smaller V1 surface areas have fewer axons and reduced orientation dispersion (left panel) relative to individuals with larger V1 surface areas (right panel).

**Supplementary Figure 2.**
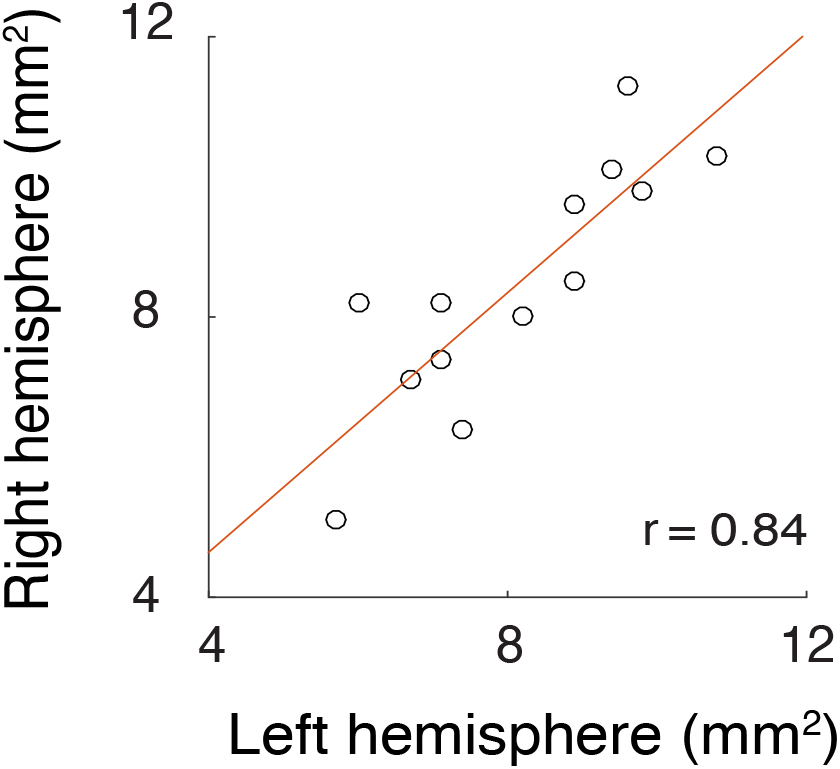
Inter-hemispheric correlation of OT cross-section area, as measured in a postmortem study (Table 2; Andrews et al., 1997).

**Supplementary Figure 3.**
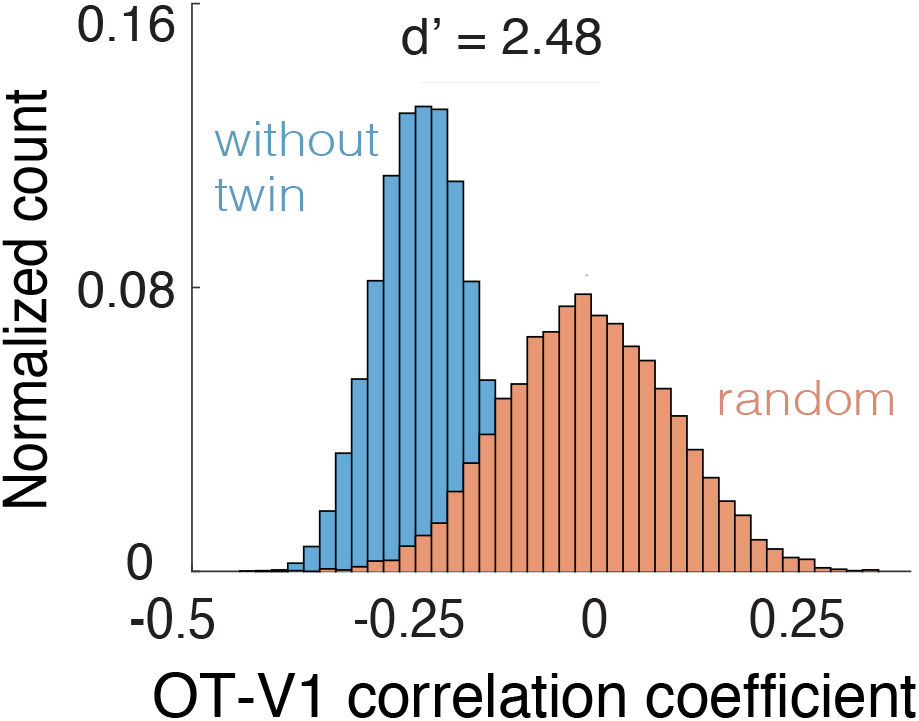
Assessment of OT-V1 correlation when excluding twin subjects. The blue histogram indicates the probability density of the correlation coefficient between V1 surface area and OT fractional anisotropy (FA) when subsampling 94 subjects data by avoiding the inclusion of both members of any twin pair. The red histogram shows the correlation coefficient when subsampling 178 subjects and shuffling the association between V1 surface area and OT FA. This histogram indicates that even if we exclude the effect of twin pairs (blue), the median correlation coefficient is almost equal to that of the main analysis (*r* = −0.21), and the distribution is clearly separate from that of random shuffling (d’ = 2.48). The vertical axis of the histogram depicts the normalized count of correlation coefficient (from all shuffles) and the horizontal axis depicts the correlation coefficient between OT FA and V1 surface area.

**Supplementary Figure 4.**
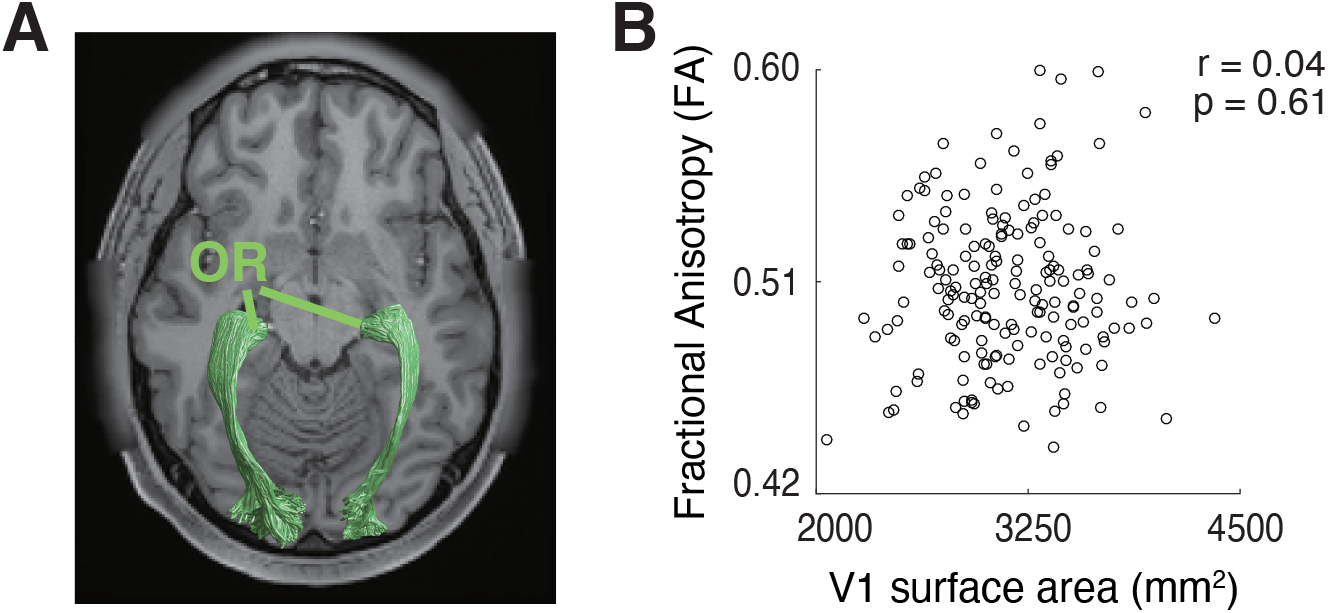
No significant correlation between V1 surface area and FA along the optic radiation (OR). **A.** The OR identified in a representative subject overlaid on an axial slice of the subject’s T1-weighted image. **B.** Scatter plot comparing V1 surface area and OR FA.

**Supplementary Figure 5.**
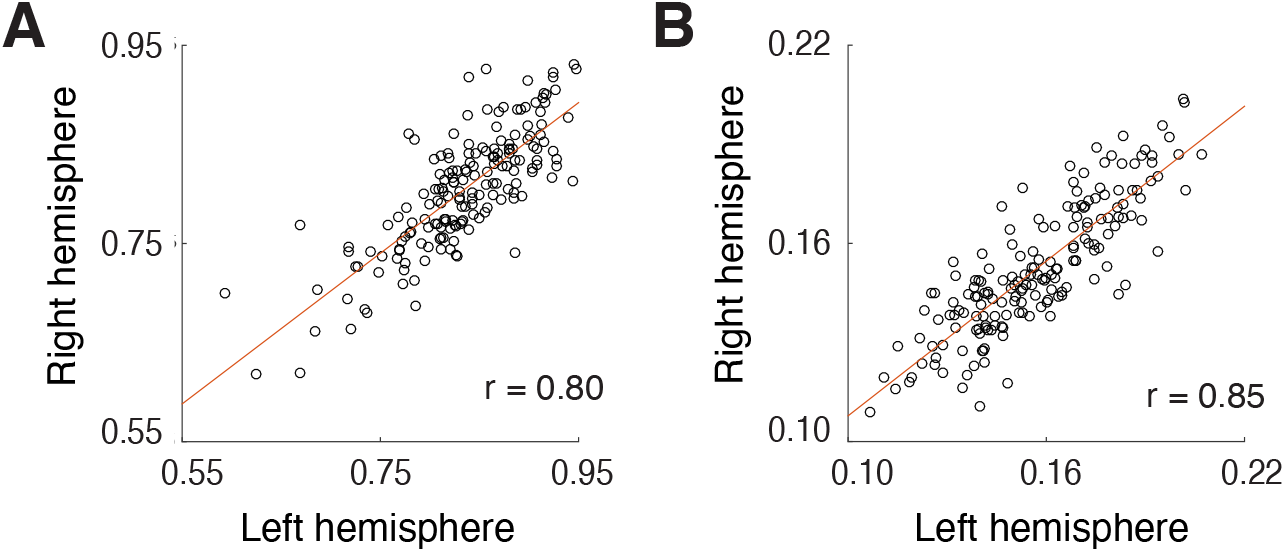
Inter-hemispheric correlation of neurite orientation dispersion and density imaging (NODDI) metrics along the OT (**A.** intra-cellular volume fraction, ICVF; **B.** orientation dispersion index, ODI). Conventions are identical to those used in Figure 2C.

**Supplementary Figure 6.**
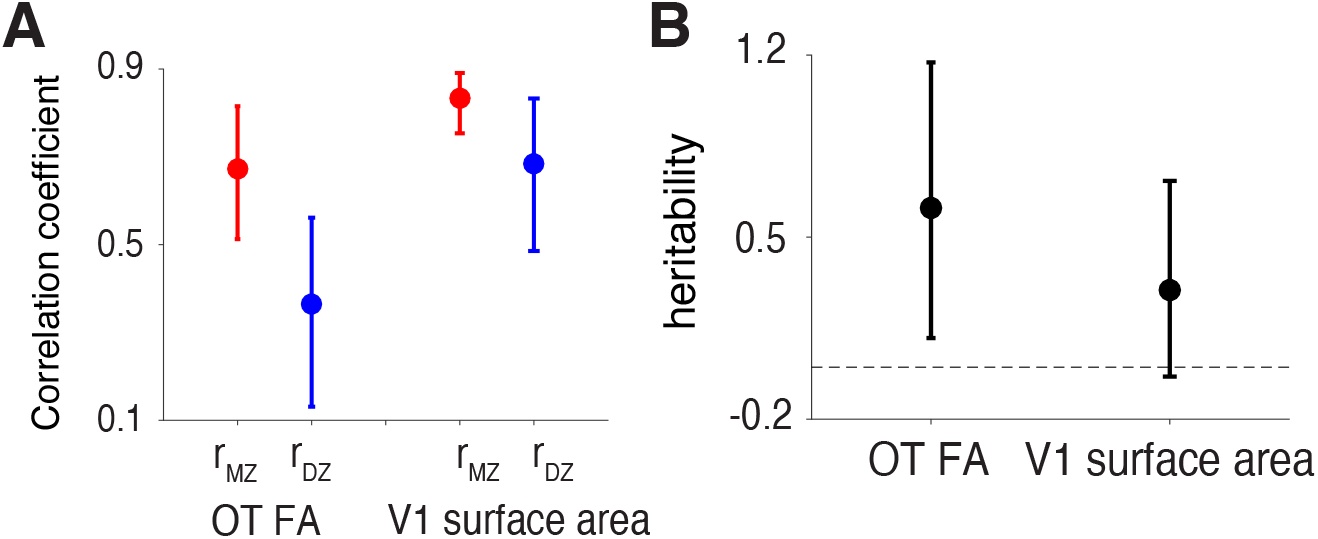
**A.** Intraclass correlation coefficient (ICC)of OT FA and V1 surface area across monozygotic (MZ, red) and dizygotic (DZ, blue) twin pairs. The error bar depicts the 95% confidence interval of the ICC estimated using 10,000 bootstraps. **B.** Falconer’s heritability index (vertical axis) of OT FA and V1 surface area. The error bar depicts the 95% confidence interval of the correlation coefficient estimated by bootstrapping.

## Notes

**Conflicts of interest:** none.

### Competing Interest Statement

The authors have declared no competing interest.

